# Re-identification of genomic data using long range familial searches

**DOI:** 10.1101/350231

**Authors:** Yaniv Erlich, Tal Shor, Shai Carmi, Itsik Pe’er

## Abstract

Consumer genomics databases reached the scale of millions of individuals. Recently, law enforcement investigators have started to exploit some of these databases to find distant familial relatives, which can lead to a complete re-identification. Here, we leveraged genomic data of 600,000 individuals tested with consumer genomics to investigate the power of such long-range familial searches. We project that half of the searches with European-descent individuals will result with a third cousin or closer match and will provide a search space small enough to permit re-identification using common demographic identifiers. Moreover, in the near future, virtually any European-descent US person could be implicated by this technique. We propose a potential mitigation strategy based on cryptographic signature that can resolve the issue and discuss policy implications to human subject research.

## Main Text

Consumer genomics has gained tremendous popularity in the last few years^1^. As of today, more than 15 million people have taken direct-to-consumer (DTC) autosomal genetic tests for self-curiosity, with about 7 million kits sold in 2017 alone^2^. Nearly all major DTC providers use dense genotyping arrays that probe ~700,000 SNPs in the genome of each participant and most DTC providers allow participants to download their raw genotype files in a textual format. This option has led to the advent of third-party services, such as DNA.Land and GEDmatch that allow participants to upload their raw genetic data in order to get further analysis (**Table 1**)^3^.

**Table 1:** Examples of vendors in consumer genetic genealogy sorted lexicographically. Database sizes were taken from ref: (*2*) based on data available on May 2018. DTC provider signifies a service that produces genomic information from biological material such as buccal swabs or saliva. Long range familial searches refer to services that employ IBD matches. 3^rd^ party support refers to services that allow upload of raw genomic data. The list includes only DTC suppliers or 3^rd^ party services with relative finder capabilities mentioned in (*2*, *3*).

| Service | Database size | DTC provider | Relative finder | 3<sup>rd</sup> party support |
| --- | --- | --- | --- | --- |
| 23andMe | 5M | • | • |  |
| Ancestry | 9M | • | • |  |
| DNA.Land | 100K |  | • | • |
| FTDNA | 1M | • | • | • |
| GEDmatch | 1M |  | • | • |
| LivingDNA | n/a | • |  |  |
| MyHeritage | 1.4M | • | • | • |

DNA matching is one of the most popular features in consumer genomics. This feature harnesses the dense autosomal genotypes to find identity-by-descent (IBD) segments, which are indicative of a shared ancestor. Previous studies have shown that this technique has virtually perfect accuracy to find close relatives and good accuracy to find distant relatives, such as 2^nd^ or 3^rd^ cousins^4–6^, providing the option for long range familial searches. This feature has led to many “success stories” by the genetic genealogy community, including the reunions of Holocaust survivors with relatives when they thought they had no living family left, reunions of adoptees with their biological families, and investigations of potential abductions of babies^7^.

From a technical and regulatory perspective, the consumer genomics tools are far more powerful for familial searches than traditional forensic techniques. Forensic familial searches use a small set of ~20 autosomal STR regions that were standardized for traditional fingerprinting^8,9^. This set is not sufficient to detect IBD matches and instead forensic techniques have to rely on partial allelic matches as a means to identify relatives. Due to the small size of the STR panel, the forensic technique is limited to finding only 1^st^ to 2^nd^ degree relatives and can suffer from false positives^9^. In addition, familial searches in forensic databases has remained a highly controversial investigative tool from an ethical and societal perspective^10^. As a result, federal legislation permits the FBI to conduct familial searches for investigating severe crimes but allows individual states to develop their own policies^11^. Some states, such as California, have placed certain restrictions on conducting familial searches, whereas other states, such as Maryland, have prohibited the practice completely. However, these limitations primarily affect searches of state-owned forensic databases and do not explicitly restrict the use of crime scene samples with civilian DNA databases, such as consumer genomics databases, yielding a much less stringent regulatory route for such searches.

In the last few months, law enforcement has started to exploit third-party consumer genetic services for long-range familial searches. This route to re-identify individuals has been predicted before^12^, but became practical only recently with the rapid increase of consumer genomics database sizes. In a recent high-profile case, the FBI used a long-range familial search to trace the Golden State Killer^13,14^. The investigators obtained a dense genome-wide genotyping profile of a crime scene sample of the perpetrator either by an array or whole genome sequencing but no cooperation of a DTC provider based on public information. They rendered the profile to look like an ordinary dataset from a consumer genomics provider and uploaded the profile to GEDmatch, a third-party website that has approximately 1 million samples that offers long range familial searches using IBD matches. The GEDmatch search identified a 3^rd^ degree cousin of the perpetrator in GEDmatch^13^. The FBI team consisted of five genealogists took four months until they were able to trace the identity of the perpetrator, which was confirmed by a standard forensic test. In a few other less notable cases over the past few months, private DNA detectives from the “DNA Doe Project” used long range familial searches with third party services to identify the remains of the bodies of “Lyle Stevik” with an estimated time of a few hundred hours of work and of the “Buckskin Girl” in a few hours of work^15^. Recently, a forensic DNA company announced that they set up a division that will use such long-range familial searches and uploaded 100 cold cases to third-party DTC services^16^. Finally, as we were authoring this manuscript, the Snohomish County Sheriff’s Office announced that they solved a cold case from 1987 of a double murder using another long-range familial search on GEDmatch^17^. In this case, the investigators found two relatives: a paternal half-first cousin, once removed and a maternal first cousin, once removed. Using genealogical searches, this profile quickly led to one sibship of three sisters and a brother. As the perpetrator was a male, the investigators focused on the brother who was confirmed to be a person of interest with a traditional forensic DNA test.

We took an empirical approach to investigate the probability of a long-range familial search to re-identify an individual. To this end, we analyzed a dataset of 600,000 individuals that were tested with a DTC provider and consented for this type of research (**Supplementary Methods**). About 85% of all individuals showed European heritage as their main DNA ethnicity (**Supplementary figure 1; Supplementary table 1**), similar to previous studies with DTC individuals^18^. We searched for IBD segments among the 180 billion potential pairs of individuals. Overall, these individuals formed a dense IBD network, with over 1 billion pairs having at least a single IBD region longer than 6cM. We derived a subset of these pairs that included only those with at least two IBD segments, which increased the chance of correctly inferring genealogical relationships. Next, we removed pairs with IBD segments greater than 700cM (approximately first cousin and closer relationships) to circumvent potential ascertainment biases due to close relatives sending in their kits together. Finally, we analyzed the number of individuals with at least one IBD segment between 30cM and 600cM (**Supplementary Methods**). The low end of our range corresponds to 4^th^ cousins; the high end corresponds to 2^nd^ cousins based on a large-scale crowd sourcing project^19^.

Our results show long range familial searches have a good probability to return a relative for a database size of 600,000 individuals (**Figure 1A**). We found that 46% of the searches will result in an IBD of at least 100cM, which typically corresponds to third cousin or closer relative, similar to the Golden State Killer Case. Interestingly, these results, which rely on a partial genetic database, are considerably higher than surname inference from Y-chromosome, which is another re-identification tactic using genetic genealogy data^20^. Moreover, long-range familial searches allow direct re-identification of females and not just males. In 10% of the searches with our data of 600,000 individuals, the top match will have an IBD segment of at least 300cM, which corresponds to a second cousin or a closer relative. To further validate our results, we also manually performed 30 random long-range familial searches in GEDmatch, which has pproximately 1M individuals in their database. The results were consistent: the top match in over 90% of he searches shared >30cM, in 75% of the searches shared >100cM match, and 10% of the searches with >300cM match (**Figure 1A**). Since most individuals in these databases are US Caucasians, these results are likely to be relevant to this ethnic group.

**Figure 1:**
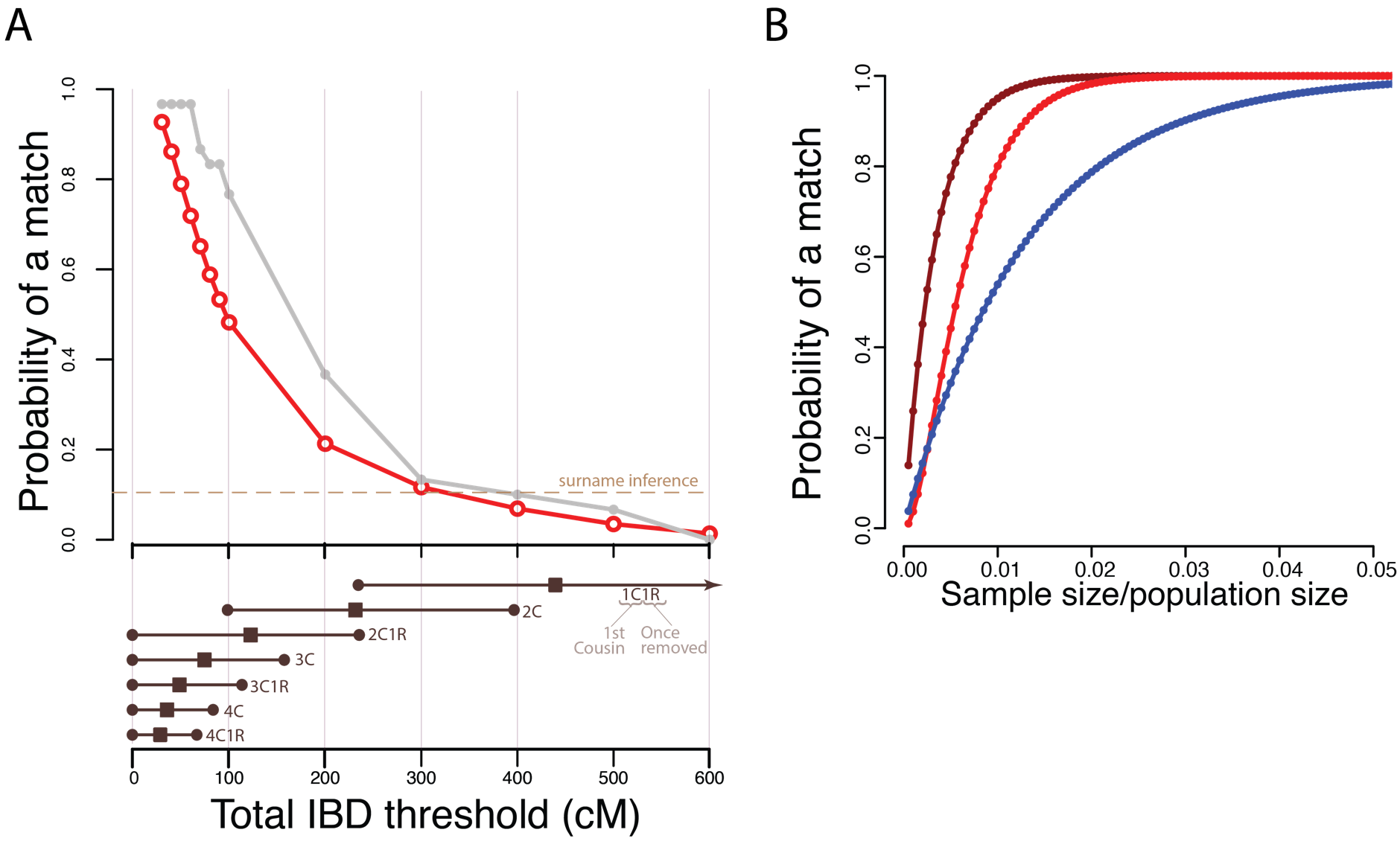
The performance of long range familial searches for various database size. **(A)** The probability to find at least one relative for various IBD thresholds (top) using 600,000 searches of DTC tested individuals and 30 random GEDmatch searches (gray). Dashed line: probability of a surname inference from Y chromosome data (21). Bottom: 95% confidence intervals (circles) and average IBD (squares) for 1^st^ cousin once removed (1C1R) to 4^th^ cousin once removed based on data in (19) **(B)** Population genetics theoretical model for the probability to find a match at least one 3^rd^ cousin or a close relative (dark red), two 3^rd^ cousins (red), or a second cousin (blue) as a function of the database coverage of the population size.

More broadly, we expect long range familial searches to return a match to virtually anyone with genetic databases that cover even a small fraction of the target population. This assertion relies on a population genetics model that takes into account the probability of sharing at least two IBD segments of >6cM and assuming population growth rates seen in the last 200 years in the Western world [a recent blog post by Doc & Coop conducted a similar analysis for GEDmatch database size^21^] (**Supplementary Methods**). This model has multiple simplifying assumptions such as no population structure, inbreeding, and random sampling of participants. However, we found that the model showed a good approximation of our empirical analysis by predicting that 44% of the searches will return at least a third cousin match compared to the observed rate of 46% for >80cM for north Europeans in our data. If considering a US Caucasian target, similar to the Golden State Killer, our model predicts that a database with ~5 million individuals (2% of this ethnic group) has a 3^rd^ cousin match for virtually any person in this ethnic group. With databases of this size, over 90% of the searches will return more than one 3^rd^ cousin, which can greatly improve triangulation and ~70% of searches will return a 2^nd^ cousin or a closer relative. Notably, consumer genomics grows at exponential rates and covering 2% of the US Caucasians is within reach for some 3^rd^ party websites in the near future.

Next, we wondered on the ability to narrow down the suspect list after finding a match in a long-range familial search. We assumed a case where a long range familial search retuned a *bona fide* match to a 3^rd^ cousin or genetically equivalent relative of >100cM. Furthermore, we considered a scenario where that the sex of the person of interest is known, their age can be estimated within a 10yr interval, and the location of residence can be estimated within a radius of 100miles (approximately the land area of the state of Maine). We used extensive genealogical records of population scale family trees^22^ to analyze whether basic demographic information has the power to quickly prune this search space (**Supplementary Methods**).

We found that the suspect list can be quickly pruned using simple demographic information. We predict that a match in the scenario has a search space of ~850 relatives on average (**Figure 2A**). Our simulations suggest that geographic data will exclude on average 57% of the list (**Figure 2B** and **Supplementary table 2**). Next, age at 10yr interval is expected to exclude another 91% of the relatives (**Figure 2C**), leading to 33 individuals on average. Finally, sex information will halve the list to around 16-17 individuals on averages, a search space that is small enough for manual inspection. In research projects, the HIPAA privacy law permits the release of the year of birth, which is even a more powerful identifier (**Figure 2D**). Our analysis shows that age at a single year resolution together with geography (<100miles) and sex is expected to return 1-2 individuals. **Figure 2E** summarizes the entire process.

**Figure 2:**
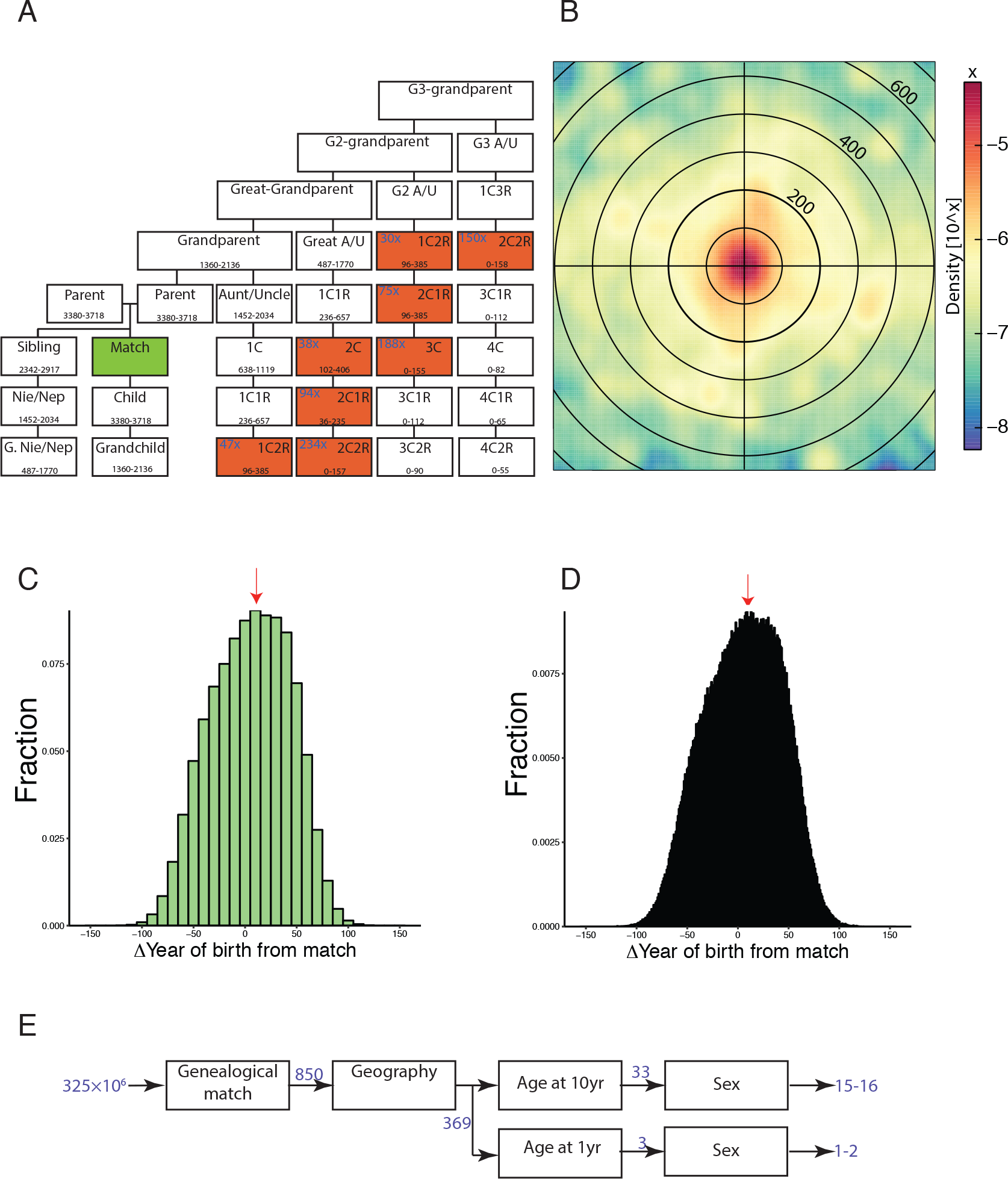
Tracing a person of interest from a long range familial match. (A) The possible relatives of a match (green) in a database. Each square represents a potential degree of relatedness. The range corresponds to the 5%-95% percentile of shared IBD in cM based ref: ^19^. Red: relatives that could fit a *bona-fida* 3C match (slightly over 100cM). The fold increases in blue denotes the average number of relatives based on a fertility rate of 2.5 children per couple. Nie/Nep: Nice/Nephew; G2: Great-great; G3: Great-great-great; A/U: Aunt/Uncle (B) An example of the geographical dispersion of 3C and 2C1R around the matched relative. Every circle denotes 100km (C-D) The distribution of the expected age differences between a match and their potential relatives with a genetic distance of third cousins. Note that some relatives of a match are yet to be born whereas other are likely to be dead. Results in the main text report the worst case scenario in which the age estimator of the person of interest is in the highest bin of each histogram (red arrow) (C) The age distribution in 10yr resolution (D) The age distribution in one-year resolution (E) The entire flow of using demographic identifiers with a long range familial match to find a US person (blue: average number of people).

Taken together, our lines of analyses show that long-range familial searches have the potential to re-identify substantial numbers of US Caucasian individuals. The main barrier is not finding a match or pruning the search space to trace the person of interest. Rather, successfully tracing an individual simply depends on the accessibility of genealogical data, their accuracy, and the determination of the investigators. Indeed, policymakers and the general public might be in favor of such enhanced forensic capabilities for solving horrendous crimes. However, we caution that the open nature of these services means that the very same technique can be exploited to identify genomic data of research subjects or counter-espionage activities by foreign adversaries.

We propose a technical measure that can mitigate some of the risks and restore control to data custodians (**Supplementary figure 2**). The collection and processing methods of traditional DTC providers are geared towards a large amount of saliva or buccal cells and not to the minute quantities of DNA and a variety of tissue of origin common to crime scene evidence. Therefore, forensic long-term familial searches have so far used special labs to develop the raw genotype data and had to render the data to mimic the format of regular DTC providers in order to be uploaded to third-party services. In our proposal, DTC providers will add cryptographic signatures to the header of the text file containing raw data available to their customers. Each supplier will use a secret private key for signing the data and will make the public key available at a known Internet address. This way, third-party services will be able to authenticate that a raw genotyping file was created by a valid DTC supplier without any modification and distinguish between valid sources and questionable sources. In case of a failure to validate the file, the 3^rd^ party service can either reject the file, allow the DNA profile to be found but not to initiate a search, or quarantine the file until some reassurance about its origin is provided. Of course, on a case-by-case basis, third-party services can cooperate with law enforcement and allow the search as opposed to the current situation in which such searches are conducted unilaterally. Similarly, this approach can also prevent exploiting long-range familial searches to re-identify research subjects.

To facilitate the adoption of our proposal, we provide a source code (under the free MIT license) that can sign and verify the raw genotype files and relies on a well-known digital signature scheme^23^. Importantly, our software does not assert or recommend any list of suppliers. Any lab that produces raw genotype files is welcome to use this scheme and any third-party provider should decide independently which list of suppliers they want to support. We believe that this technical approach, if adopted, can quickly mitigate some of the risks compared to legislation that usually takes a considerable amount of time^24^.

The rise of long-range familial searches also has implications for human subject research. The U.S. Department of Health and Human has recently rejected proposals to include whole genome sequencing as identifiable information in the Revised Common Rule but implemented a mechanism to evaluate the scope of identifiable private information based on new technological developments^25^. The growing success of long-range familial searches shows that even simpler types of genotypic information, such as genome-wide genotyping arrays, can be used to identify individuals with high success rates. These rates will grow in the near future due to the immense interest in consumer genomics. These developments will further challenge the status quo regarding the identifiability of DNA data of human subjects and may require the developments of new policy measures to further protect these datasets.

## Acknowledgments

We thank G. Japhet for his contributions to the cryptographic signature scheme and Y. Naveh for valuable comments. Y.E. holds a Burroughs Wellcome Fund Career Awards at the Scientific Interface. Y.E. conceived the idea for this study. Y.E. and T.S conducted the analysis of matches using the MyHeritage and the Geni.com data. S.C and I.P. developed the theoretical framework to estimate the number of matches. Y.E. and T.S. adapted the code for the cryptographic signatures. Y.E., T.S., S.C., and I.P. wrote the manuscript. The code for the cryptographic signatures is available on https://github.com/erlichya/signature.

## Supplementary Methods

### 1. Measure shared IBD with a subset of the MyHeritage database

The MyHeritage database mainly consists of individuals that were tested with the MyHeritage DNA product. Briefly, individuals swab the inner side of their cheeks using a sterile absorbent tipped applicator (HydraFlock). After sampling DNA on the inner side of the cheeks, the participant places the tip of the applicator in a vial that is filled with a standard lysis buffer. The DNA is transferred to a CLIA certified lab, where is genotyped with an Illumina OmniExpress genome-wide genotyping array that contains 700,000 SNPs. Another route for participants to enter the database is by uploading their raw genotype files from other DTC companies. Currently, the website supports uploads from 23andMe (v1-v4 kits), Ancestry (all versions), and FTDNA (all versions). All participants have agreed to the MyHeritage’s Terms and Conditions that permits genetic analysis of their data.

To measure the probability of finding a relative above a certain shared IBD, we took the results from the standard DNA processing pipeline of MyHeritage, which lists all IBD segments above 6cM for pairs of individuals. For this study, we used 600,000 samples, which is a subset of the MyHeritage database. The IBD segments of this subset of samples are stored in a special research database in a de-identified format that is capable of fast computing and represent the DNA data the company had in the summer of 2017.

Next, we queried the database with various levels of minimal shared cM between the relatives. In our experience, customers tend to purchase more than one kit and hand the other kit to a close family member. To mitigate ascertainment bias, we deliberately excluded all pairs of individuals above 700cM that are likely to be first cousins or closer relatives. As the service also offers individuals to document their family trees, it is often possible to find known genealogical relationships between IBD matches. To further reduce potential ascertainment biases, we excluded pairs of relatives with known genealogical paths of up to first cousins or relationships with similar genetic distance such as grandparents. Finally, we queried the database with thresholds growing from 30cM to 600cM and counted the number of individuals with at least one match.

To calculate the genetic ethnicity of each user, we used the standard results of the MyHeritage ethnicity pipeline. This pipeline reports 42 possible ethnicities based on a reference dataset of over 5000 samples collected from MH participants that consented for this process and presented homogenous ethnicity as reported by the place of births of their ancestors. For the purpose of this study, we assigned each ethnicity to eight coarse subcontinental regions (**Supplementary table 2**).

### 2. Measure shared IBD with the GEDmatch database

GEDmatch employs a unique model where each user is allowed to search any kit number in their database using the “One-to-many DNA comparison” tool. Importantly, the raw DNA data of the kit is not available to the user. However, the website does allow users to view a list of matches of any other kit that opted-in to the “One-to-many DNA comparison” tool. The website sorts the list of matches by the total shared IBD and includes key details about the match including the contact information of the user that uploaded the matched kit and in some cases also pedigree information.

We manually viewed the match lists of 30 kit numbers using the “One-to-many DNA comparison” tool. We selected the default parameters that requires IBD matches to be at least 7cM long. The kit numbers were selected by a random number generator to avoid potential biases. For each match list, we examined the total IBD of the top match. If the match was over 700cM, we excluded it from the list, similar to the exclusion process with the MyHeritage data described above. Finally, we examined the top 30 results and filtered the list according to various cM thresholds, which gave the results reported in the main text.

### 3. Population genetic theoretic calculation of long range familial searches

#### The problem

Consider a database of genotyped individuals from a defined population and the DNA of a person of interest, called the target. We would like to identify the target by finding his/her relatives in the database. We wonder about the probability to find a relative given the population size, the database size, and the matching parameters.

#### The model

- We assume a monogamous Wright-Fisher model, similar to that of Shchur and Nielsen ^26^. In the current generation, the population has ***N*** males and ***N*** females, organized in ***N*** mating pairs (couples). Each individual in the current generation chooses its parents (i.e., a mating pair) randomly out of all mating pairs in the previous generation.
- The number of children per mating pair is ***r* > 2**. Thus, the population size at generation *g* before the present is ***N*(*g*) = *N*(*r*/2)^−*g*^**.
- Individuals are diploid and we consider only the autosomal genome.
- The database has ***R*** individuals.
- The target individual is compared to all individuals in the database. We only consider relationships up to ***g*_max_**-cousins. For example, 1-cousins are siblings, 2-cousins are first cousins, etc. All cousins/siblings are full.
- If the target is a degree ***g* ≤ *g*_max_** relative of an individual in the database, their (diploid) genomes are compared, and identical-by-descent (IBD) segments are identified. We assume that detectable segments must be of length **≥ *m*** (in Morgans) and that to confidently detect the relationship (a “match“), we must observe at least ***s*** such segments.
- The number of matches between the target and the individuals in database is counted. If we have more than ***t*** matches, we declare that the target individual has been “identified”. Typically, we simply assume t=1, as in the Golden State Killer case.

#### Derivation

##### The probability of a shared mating pair between the target and a single reference sample

Consider two individuals: the target and a single individual in the database. ***g*** generations before the present, each one of them has **2**^***g*−1**^ ancestral mating pairs (containing **2**^***g***^ ancestral individuals). For example, each individual has one pair of parents (***g*** = **1**) and two pairs of grandparents (***g*** = **2**). For large ***N***(***g***) (**2**^***g***^ ≪ ***N*(*g*)**), the probability that the two individuals share an ancestral mating pair is approximated by:

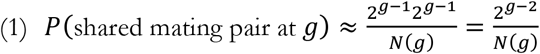

Each ancestral mating pair of the target has a probability of **1 − 2^*g*−1^/*N*(*g*)** not to share any ancestor with the database individual. The probability of all ancestral mating pairs of the target are not shared with the reference is 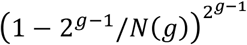, and the probability that at least one is shared is 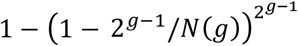. Eq. (1) is the limit when **2^*g*−1^ ≪ *N*(*g*)**. We ignore the possibility of sharing more than one ancestral mating pair, assuming, similarly, that **2^2*g*^ ≪ *N*(*g*)**.

The probability to share an ancestral mating-pair for the first time at generation *g* is approximately

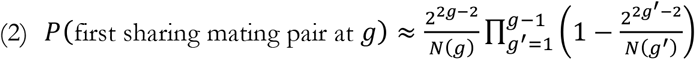

##### The probability of a match given a shared mating pair

Next, we determine the probability that the target and his/her relative via ancestral mating pair g generation above are identified as a match using genetic data. To this end, we calculate the probability that they share at least *s* IBD segments longer than *m*.

We use a simple approximation that the genome can be broken into effectively independent blocks, each of which is inherited independently. If the ancestors have lived ***g*** generations ago, the two individuals are separated by **2*g*** meioses. Given that the total genome length is roughly ***L* = 35**Morgan, there are on average **2*gL* ≈ 70*g*** recombination events between the two individuals. Since blocks can also be bounded by chromosome ends, a rough approximation for the number of blocks is **2*Lg* + 22**, as in ref. ^27^.

In each block, a pair of individuals share one ancestral mating pair out of **2^*g*−1^**. The probability that the relevant lineage in each individual descends from the shared mating pair is **1/2^*g*−2^** (for ***g* > 1**), leading to a probability of **1/2^2*g*−4^** that both lineages descend from the shared mating pair. Note that this is taking into account the fact that we can detect IBD sharing between any of the two chromosomes in one individual and any of the two chromosomes in the other. Thus, we can go “up the tree” in the right direction towards the shared ancestors with a probability of 1.

The individuals in the shared mating pair have four chromosomes in total. For any one of them chosen by the target, it will be chosen by the reference with probability **1/4**. Thus, the probability of sharing the same chromosome is **1/4** ‧ **1/2^2*g*−4^ = 1/2^2*g*−2^**. For ***g* = 1**, the probability comes out as **1**, which is a reasonable approximation given that full siblings share one chromosome in about half of their genomes, and two chromosomes in a quarter of it.

Next, we determine that probability of the IBD segment to be sufficiently long given that the pair share an ancestral chromosome. The length of the segment is exponential with rate **2*g*** (with length measured in Morgans), namely, if *x* is the segment length, ***P*(*x*) = 2*ge*−^2*gx*^**. The probability of the segment length to exceed ***m*** is 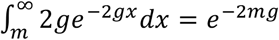. Thus, in each block, the probability of sharing a detectable IBD segment is

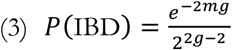

Assuming that blocks are independent, the probability to share ***k*** blocks is binomial with ***n* = 2*Lg* + 22** (the number of blocks) and ***p* = *P*(IBD)** above. To declare a match, we need at least ***g*** segments. Thus, given a shared mating pair *g* generations ago, the probability to observe a match is

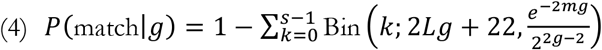

##### The number of matches to the database

We approximate the probability of a match by assuming that the set of ancestors for all individuals in the database are non-overlapping. This assumption was recently demonstrated by Shchur and Nielsen^26^ for a constant population when ***N* → ∞** and **R/N** is constant. Thus, the probability of a match with one specific individual in the database can be obtained simply by summing Eqs. (2) and (4) over all *g*,

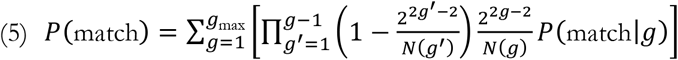

Finally, to identify an individual, we need to find at least t matches in the database. If matching is approximately independent across the individuals in the database, then the probability of identification is binomial, with ***n* = *R*** and ***p* = *P***(match) from Eq. (5). Thus,

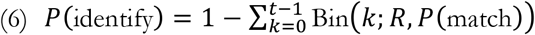

Eq. (6) is our final expression.

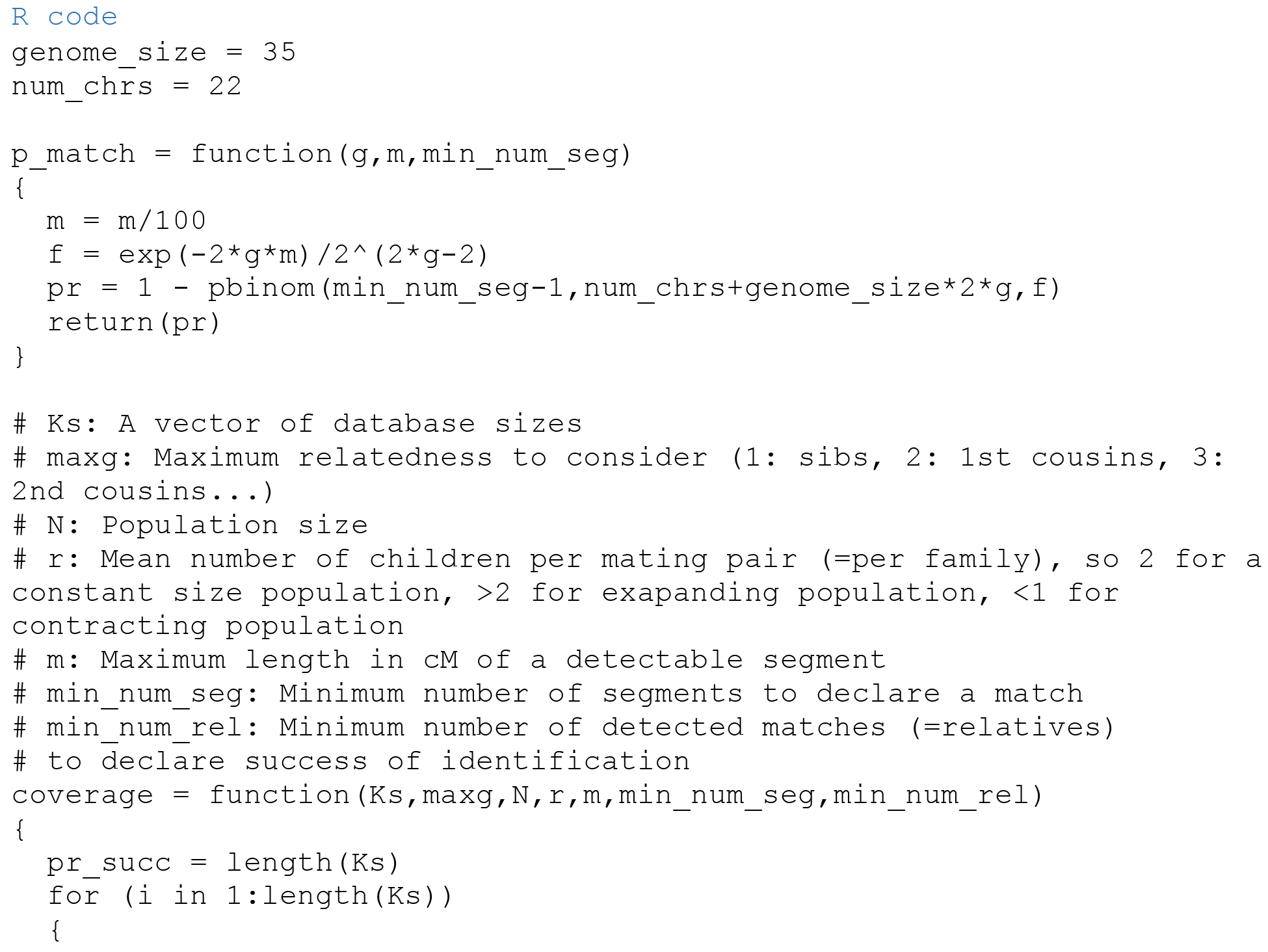
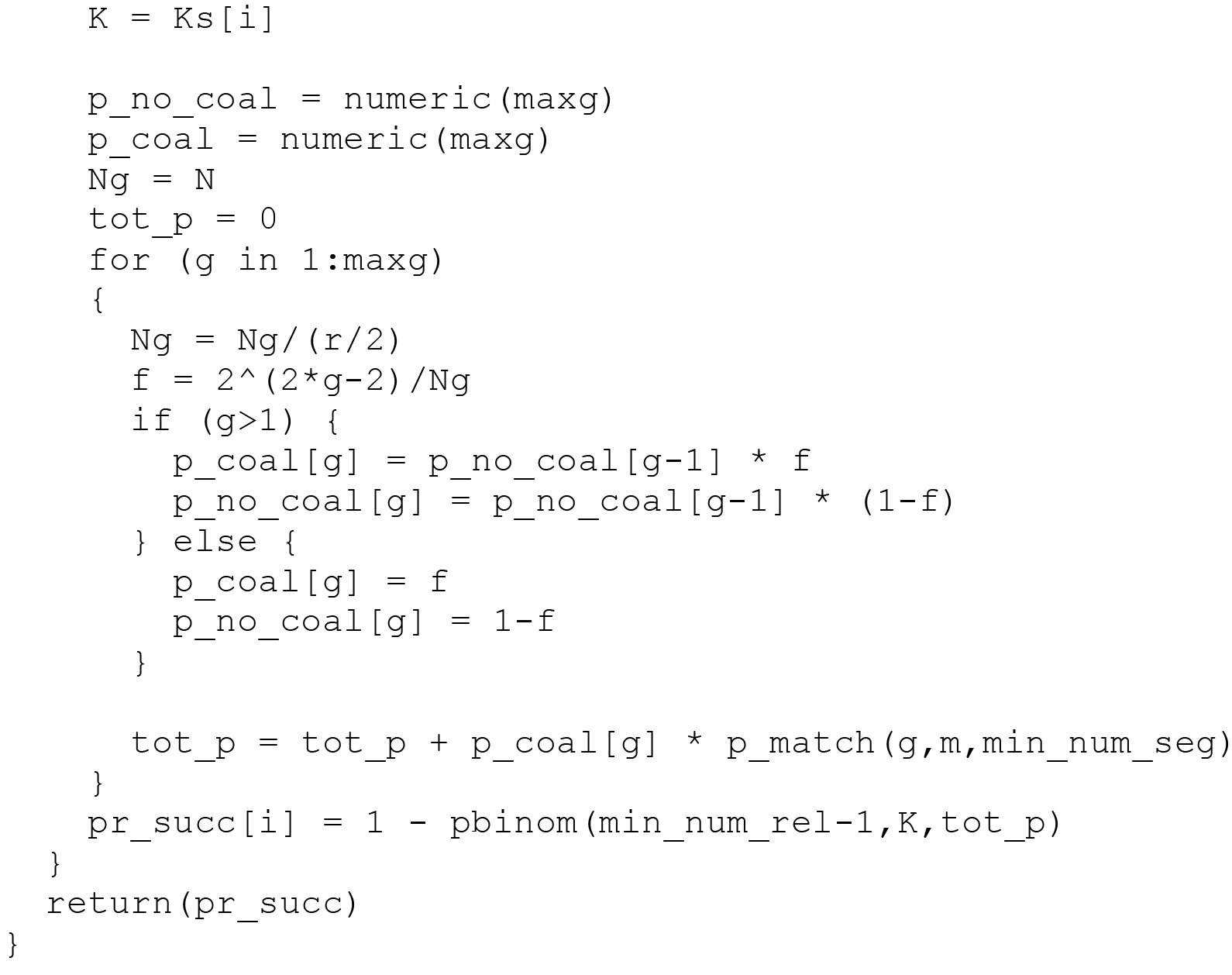

To produce Figure 1B, we the following parameters with the **R** code above:

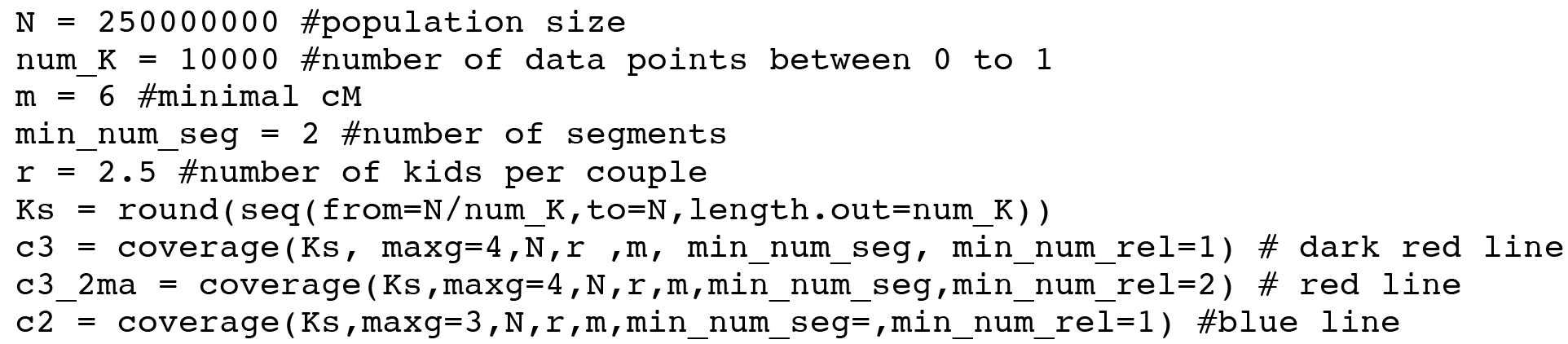

### 4. Pruning the search space with genealogical identifiers

To find the power of demographic information to prune the search space, we turned to our previous study with population scale family trees ^22^. This dataset represents a collection of large scale family trees and has been subject to extensive types of validation, including accuracy assessment using genetic data and concordance analysis with government-based demographic data.

We focused on the following types of relatives: 1C2R, 2C1R, 3C, 2C, and 2C2R types as these genealogical relationships are typical for genetic distances of ~100cM. We did not take into account rarer types of pairs with similar genetic distance such as half-second cousins as these are less likely to be encountered. To estimate the number of individuals in each class, we assumed that each couple gives birth to 2.5 kids on average and that all kids reaches a fertility age similar to ref. ^6^. In total, we estimated that 855 relatives exist with a genetic distance of slightly over ~100cM.

#### Geography

We used our genealogical records to analyze the geographic distance between relatives. We analyzed 145,658 pairs of relatives encompassing 1C2R, 2C1R, 3C, 2C, and 2C2R (**Table S2**). We considered only pairs where at least one individual was born in the US between 1940 to 2010. We then calculated the geographical distance based on the longitudes and latitudes of their birth locations.

We consider a conservative scenario in which the potential residence of the person of interest centers around the matched relative. When the search radius is restricted to 100miles from the match, less than 30% of all matching 1^st^ cousins once removed are expected to within the search space centered around the match, whereas 51% of all matching 2^nd^ cousins are expected to be around this search space.

After considering the number of relatives for each class and their geographical distribution, our model estimates that on average only 369 relatives live within 100 miles from the match.

#### Age

To analyze the age dispersion of pairs of relatives, we conducted extensive simulations that were further validated with a large set of 3^rd^ cousins.

##### Simulations

Genealogically, the year of birth differences between various types of cousins is expressed as three processes: (i) the year of birth differences of two siblings, denoted by s (ii) the sum of the parent age at birth of *i* consecutive descendants of one sib, denoted by v_i_ (iii) the sum of the parent age at birth of j consecutive descendants of the other sib, denoted by u_j_. For example, for 2^nd^ cousin once removed (2C1R), the difference is s+v_3_−u_2_. In general, for a x-cousin y-removed pair, the difference in the year of birth is s+v_x_−u_(x-y)_.

To simulate s, v, and u, we examined the parent age at birth of 1.74 million parent-offspring pairs that reflect the highest quality of our data with exact date of births and birth places. Next, we created a histogram of the differences and excluded a handful of event with birth year difference of less than 10 years or more than 60 years that is likely to stem from typos of genealogists. We created a similar histogram for 870,000 pairs of full siblings. To simulate an instance of v_x_ or u_x_, we randomly sampled *x* events according to the probability mass function of the parent-age histogram and summed them together. To simulate an instance of s, we sampled an event from the probability mass function of the sibling year of birth difference histogram.

We simulated 100,000 cases of each type of relative of an interest. When simulating these pairs, we took into account interdependencies between the generations. For example, the age difference of 1C2R was the starting point for the age difference of 2C1R, which then was the starting point of the age difference of 3C. We then calculated the density of age distribution of each class and mix them according to the number of expected people in each class times the probability of a person living less than 100miles from the match. The entropy of the histogram at 10yr bins was 3.955bits and the entropy of the 1yr histogram was 7.26bits (entropy is a good measure for identifiability because it describes the expected information content of a piece of information).

In the highly conservative scenario, the age of the person of interest would fall under the highest bin of the histogram. We measured this bin for the 10yr interval and 1yr interval and reported the results in the main text.

##### Direct analysis of a large set of relatives

Our data also allows to measure the year of birth differences in a large set of known distant relatives as the geography analysis above. However, age analysis is more complicated when measured with recently born relatives due to ascertainment bias issues. First, the simulations above showed that for some types of relatives such as third cousins, the potential relative can be 70 years younger than the examined person, meaning that the relative is yet to be born, creating a censoring effect on our data. Second, in our previous studies with this dataset, we find that most individuals in our data came from the late 19^th^ century due to the tendency of genealogists to document ancestors of their families. We were concerned that relatives ascertained from recently born individuals would disproportionally reflect these old cases and skew the age analysis.

As an alternative, we focused on historical data rather than recent data. We ascertained 1.2 million pairs of 2^nd^ cousins and 1.7 million pairs of 3^rd^ cousins that were born between 1800 to 1910. All of these pairs had exact birth date data and known birth locations. We found that the differences in the year of the actual 2^nd^ and 3^rd^ cousins were relatively similar to their simulations:

|  | Entropy of<br>observed data<br>[bits] | Entropy of<br>simulation [bits] |
| --- | --- | --- |
| 2C – 1yr resolution | 6.04 | 6.17 |
| 2C – 10yr resolution | 2.74 | 2.87 |
| 3C – 1yr resolution | 6.25 | 6.40 |
| 3C – 10yr resolution | 2.95 | 3.09 |

These small differences can stem from other types of ascertainment biases with the historical data or the resemblance between relatives that induces reproduction at similar ages. Nevertheless, the overall consistency indicates that the simulation captures the overall distribution of ages in this class of relatives.

## Supplementary Figures

**Supplementary figure 1:**
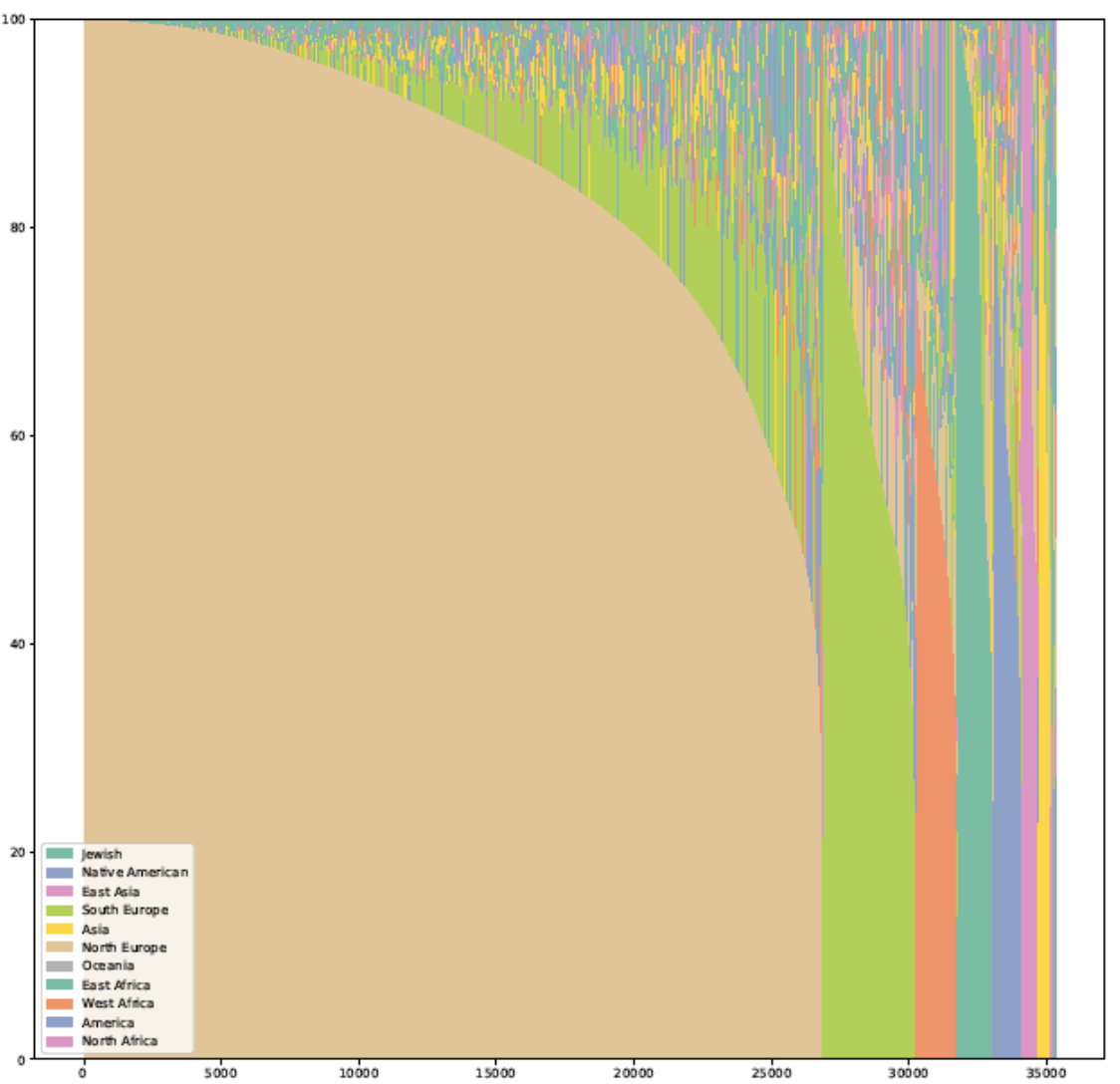
The genetic ethnicity of a 36000 users in the matching database. Each vertical line corresponds to a person and the Y-axis reflects the ethnicity composition from 0 to 100%. Colors denote the main ethnic groups in this analysis. The individuals are sorted based on their inferred main ethnicity.

**Supplementary figure 2:**
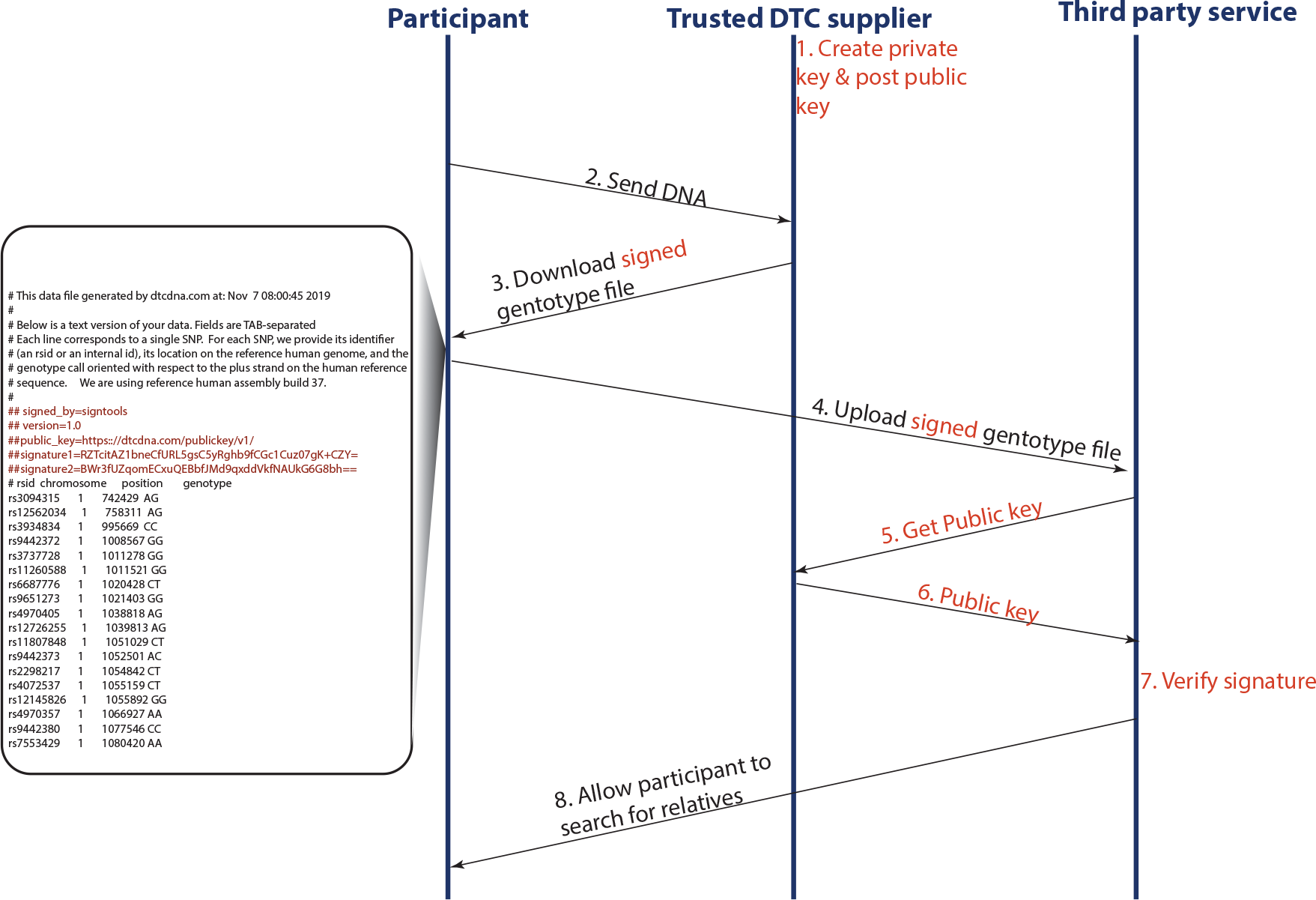
The flow of data for validating files. The chart shows how third party services can work together with trusted DTC suppliers in order to validate the authenticity of the data. Black: current flow of information. Red: added steps to authenticate the file. On the left a snippet of a raw genotype file after signing.

**Supplementary Table 1:** The fraction of individuals in each major genetic ethnicity in the 600,000 individuals in our dataset.

| Main DNA ethnicity | Percentage |
| --- | --- |
| North Europe | 76.03% |
| South Europe | 9.55% |
| West Africa | 4.39% |
| Ashkenazi Jewish | 3.59% |
| Native American | 3.03% |
| East Asia | 1.71% |
| South/West Asia | 1.29% |
| North Africa | 0.20% |
| Oceania | 0.14% |
| East Africa | 0.05% |

## Supplementary Tables

**Supplementary table 2:** The probability that a relative is found outside of a 100km,100 miles, and 200km range from a match.

| Geography | 1C2E | 2C1R | 3C | 2C2R | 2C |
| --- | --- | --- | --- | --- | --- |
| #cases | 33974 | 32432 | 13522 | 58018 | 7712 |
| >100km | 0.679049 | 0.627467 | 0.606567 | 0.664897 | 0.563797 |
| >100miles | 0.5994 | 0.539097 | 0.543854 | 0.594746 | 0.489627 |
| >200km | 0.566021 | 0.508263 | 0.522556 | 0.570616 | 0.451504 |

